# Cellular Calcium Activity at Depth Predicted from Surface Potential Recordings using Ultra-high Density Transparent Graphene Arrays

**DOI:** 10.1101/2023.10.05.561133

**Authors:** Mehrdad Ramezani, Jeong-Hoon Kim, Xin Liu, Chi Ren, Abdullah Alothman, Chawina De-Eknamkul, Madison N. Wilson, Ertugrul Cubukcu, Vikash Gilja, Takaki Komiyama, Duygu Kuzum

**Author notes:** These first authors equally contributed to this study (Mehrdad Ramezani, Jeong-Hoon Kim). **Correspondence:** Dr. Duygu Kuzum Electrical & Computer Engineering University of California San Diego 9500 Gilman Drive MC 0407 La Jolla, CA, USA 92093.

## Abstract

Recording brain activity with high spatial and high temporal resolution across deeper layers of cortex has been a long-sought methodology to study how neural information is coded, stored, and processed by neural circuits and how it leads to cognition and behavior. Electrical and optical neural recording technologies have been the key tools in neurophysiology studies toward a comprehensive understanding of the neural dynamics. The advent of optically transparent neural microelectrodes has facilitated multimodal experiments combining simultaneous electrophysiological recordings from the brain surface with optical imaging and stimulation of neural activity. A remaining challenge is to scale down electrode dimensions to single -cell size and increase the density to record neural activity with high spatial resolution across large areas to capture nonlinear neural dynamics at multiple spatial and temporal scales. Here, we developed microfabrication techniques to create transparent graphene microelectrodes with ultra-small openings and a large, completely transparent recording area. We achieved this by using long graphene microwires without any gold extensions in the field of view. To overcome the quantum capacitance limit of graphene and scale down the microelectrode diameter to 20 μm, we used Pt nanoparticles. To prevent open circuit failure due to defects and disconnections in long graphene wires, we employed interlayer doped double layer graphene (id-DLG) and demonstrated cm-scale long transparent graphene wires with microscale width and low resistance. Combining these two advances, we fabricated high-density microelectrode arrays up to 256 channels. We conducted multimodal experiments, combining recordings of cortical potentials with high-density transparent arrays with two-photon calcium imaging from layer 1 (L1) and layer 2/3 (L2/3) of the V1 area of mouse visual cortex. High-density recordings showed that the visual evoked responses are more spatially localized for high-frequency bands, particularly for the multi-unit activity (MUA) band. The MUA power was found to be strongly correlated with the cellular calcium activity. Leveraging this strong correlation, we applied dimensionality reduction techniques and neural networks to demonstrate that single-cell (L2/3) and average (L1 and L2/3) calcium activities can be decoded from surface potentials recorded by high-density transparent graphene arrays. Our high-density transparent graphene electrodes, in combination with multimodal experiments and computational methods, could lead to the development of minimally invasive neural interfaces capable of recording neural activity from deeper layers without requiring depth electrodes that cause damage to the tissue. This could potentially improve brain computer interfaces and enable less invasive treatments for neurological disorders.

## Introduction

Understanding complex dynamics of the brain and the central nervous system require studying mechanisms and functions in a diverse set of spatial and temporal scales [1–3]. Spatial scales encompass neural circuits in millimeters or centimeters, single neurons in microns, synapses in submicrons and proteins such as ion channels and receptors at the nanoscale. This spatial diversity also cultivates temporal diversity where some molecular processes are taking place in microseconds, action potentials in miliseconds, neurotransmitter or hormone release in minutes and learning and behavioral changes in hours to days [1]. Monitoring neural dynamics and interrogating neural function across these diverse spatial and temporal scales is not possible using a single tool or technology. Therefore, integration of multiple tools and sensing and stimulation modalities in the same experiment have been widely employed to link mechanisms and functions operating at these different spatiotemporal scales towards a more comprehensive understanding of the brain.

To date, multimodal experiments have been used to investigate the neural dynamics with applications ranging from studies of neural circuits [4–10] or pathophysiology of brain disorders such as Parkinson’s disease [11], Alzheimer’s disease [12], and Schizophrenia [13–17] to hybrid brain computer interfaces (BCI) combining two different modalities with complementary strengths to enhance performance [18–21]. Among these multimodal approaches, experiments concurrently recording electrophysiological during optical imaging and optogenetic stimulation has become a powerful approach to (i) combine temporal resolution advantage of electrophysiology with high spatial resolution and cell-type specificity of optical methods, (ii) to bridge the knowledge gap between basic neuroscience research relying on optical methods employing genetic modifications and clinical research mainly using electrical recordings, and (iii) to expand spatial reach of neural recordings [22] and (iv) to identify cell types through opto-tagging during electrophysiological recordings of neuronal spikes [23, 24]. To enable crosstalk and artifact free integration of electrical and optical modalities, transparent graphene electrodes with artifact-free recording capability have been proposed [25–29]. Among all materials, graphene provides the best of both worlds by combining transparency, artifact-free recording capability [25, 26], flexibility [30], low noise [31], biocompatibility [32, 33], and chronic reliability [34–36]. Other materials have also been investigated to fabricate transparent electrodes, such as indium-tin-oxide (ITO) [37, 38], carbon nanotube meshes (CNTs) [39], metal nanowires, meshes or grids [40–43], and PEDOT:PSS [44–46]. However, several constraints limit the use of these materials as multimodal chronic interfaces. The brittle nature of ITO makes it susceptible to crack formation and mechanical degradation [47, 48]. CNTs and nanowires have shown cytotoxicity in many studies raising concerns on biocompatibility [49, 50]. Metal nanowires and meshes might still absorb light leading to light induced artifacts in electrical recordings due to photovoltaic and photothermal effects [44, 51, 52]. PEDOT:PSS might exhibit chronic reliability issues due to delamination [53]. Transparent graphene electrodes have been successfully employed in multimodal studies previously [22, 25–27, 35, 54–60]. However, all graphene arrays demonstrated to date had low channel counts (∼16) and large electrode opening sizes (∼50 um or larger) limiting the spatiotemporal resolution of recorded signals. Reducing electrode dimensions to single cell size is desirable to detect high frequency activity including multiunit (MUA) and single unit (SUA) activities with high signal to noise ratio [61]. Increasing the array density and channel count is necessary to capture neural dynamics with high spatial resolution across large areas [62, 63]. Two important challenges remain to be addressed to realize high-density transparent graphene arrays with ultra-small electrodes: (1) In order to keep the field of view (FoV) clear, microwires of the arrays need to be completely transparent, particularly for high-density arrays. That requires patterning thin and long graphene wires. However, that results in increased wire resistance leading to signal attenuation and increases the susceptibility to structural defects from growth or fabrication causing open circuit failures. (2) Scaling down the graphene electrode dimensions drastically increases the impedance due to the quantum capacitance [56], an intrinsic property of graphene due to its unique band structure [64].

In this work, we overcome these challenges and demonstrate completely transparent, high-density, microelectrode arrays with ultra-small graphene electrodes for multimodal experiments. We reduced the sheet resistance of graphene wires 7-fold by adopting double layer graphene and interlayer nitric acid doping and realized high aspect ratio graphene wires with high yield. We fabricated high-density graphene arrays up to 256-channels without any metal wires in the field of view to prevent any shadows that can block the imaging field of view and to cause light-induced artifacts. To overcome the quantum capacitance and lower the impedance of small graphene electrodes, we employed platinum nanoparticles (PtNPs) and achieved low impedances (∼300 kΩ) for electrodes with 20 μm diameter. We implanted these transparent, high-density electrodes over the visual cortex of awake mice and performed simultaneous two-photon calcium imaging at different depths. These experiments enabled simultaneous recordings of cortical potentials and neural activity from multiple cortical layers, which provide mutual and modality-specific information on neural dynamics. Our analysis showed that the surface potentials at high frequencies are highly correlated with the average calcium activities of L2/3 neurons. We trained recurrent neural networks (RNNs) using the multimodal dataset acquired from these experiments to predict the average calcium activities at L1 and L2/3 from surface recordings. Moreover, we extracted a representative latent space from the neural population’s calcium response and trained RNNs to decode the latent variables (n=8). Next, we projected the decoded latents back to the original space to predict single-cell activities of L2/3 neurons. Our results demonstrated that the average (L1 and L2/3) and single-cell (L2/3) calcium activities could be predicted from the surface potentials recorded by our graphene electrodes.

## Results

### Elimination of defects and reduction of resistivity to enable large area high-density transparent arrays

To build large area and high-density graphene arrays with a completely transparent recording area, we needed to reduce the width of the graphene wires to increase the electrode density without causing a huge increase in wire resistance. Conventional metal microwires can offer low resistivity. However, they completely block the field of view for high-density arrays, which would prevent multimodal imaging (**Figure S1a**). Previous designs of transparent graphene arrays using monolayer graphene required Au wires surrounding the recording electrode area, which limited the field of view and increased the potential for light-induced artifacts [26]. Therefore, microwires of the high-density arrays have to be made of graphene to maintain complete transparency across the entire recording area (**Figure S1b**). Compared to conventional metal microwires with finite thicknesses, graphene has relatively high sheet resistance due to its single-plane 2D atomic structure and grain boundaries. Therefore, reducing the width and increasing the length of graphene wires can significantly increase the wire resistance and lead to attenuation of the recorded signals. Furthermore, thin and long graphene wires are susceptible to defects from the growth and fabrication processes. These defects increase the probability of having open circuits in the graphene wires and reduce the yield of the graphene microelectrode array.

Here, we addressed these challenges by introducing interlayer-doped double layer graphene (id-DLG) to build flexible and transparent arrays with low resistance long graphene wires and ultra-small microelectrodes (**Figure 1a)**. To form id-DLG layers, the first graphene layer was transferred with the electrochemical delamination transfer method and doped by dipping it in a 50% nitric acid (HNO_3_) solution. Then, the second graphene layer was transferred using the same method as the first graphene layer to cap the interlayer dopants, as shown in **Figure 1b**. Trapping dopants between two graphene layers is important for achieving stable doping and corresponding resistivity decrease in graphene layers [65]. Details of the fabrication steps are explained in the **Methods** section.

**Figure 1.**
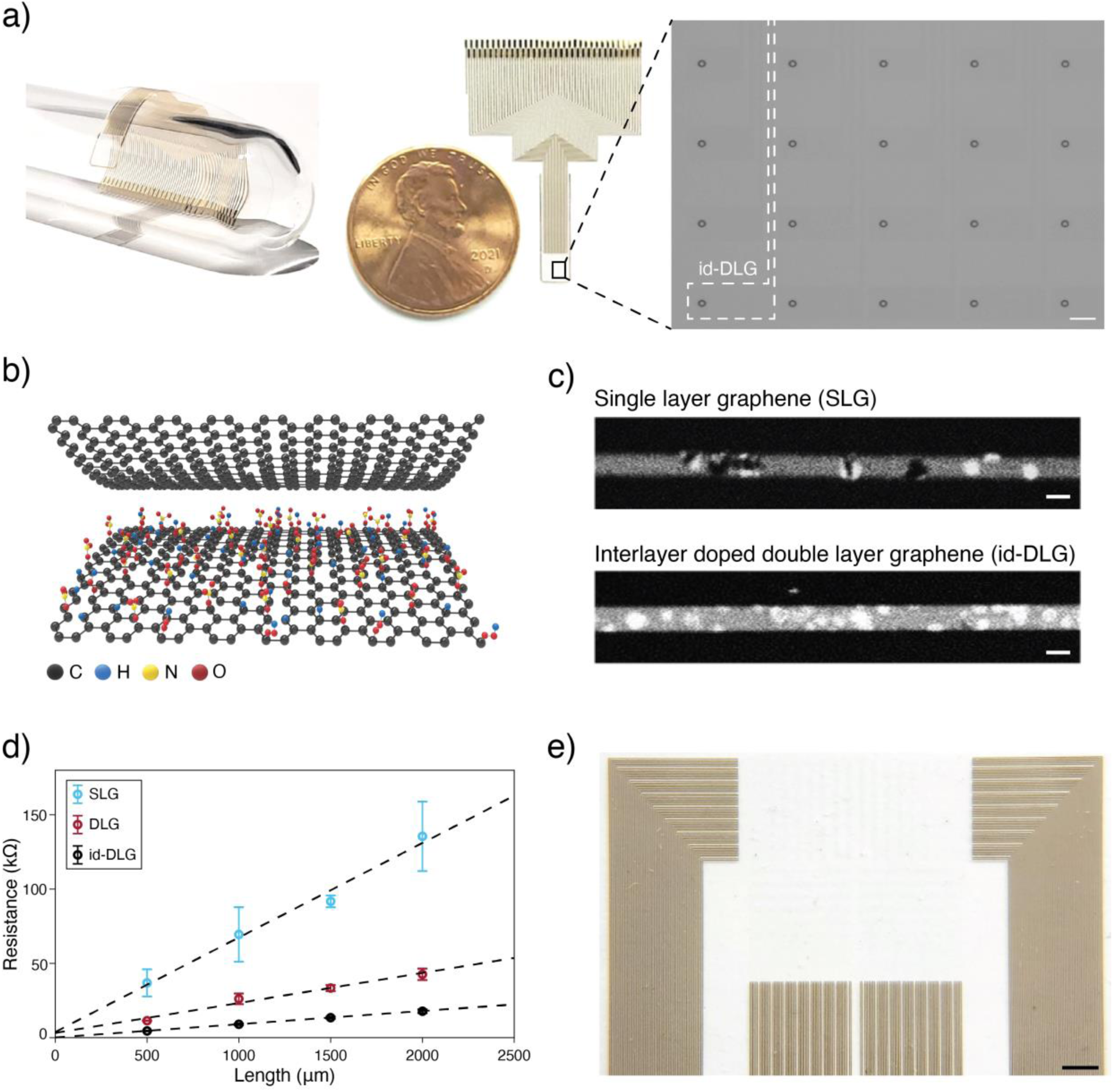
High-density transparent graphene array. (**a**) Transparent and flexible 64-channel graphene array (left) and magnified part of it with graphene wires shown with white dashed lines (right). The scale bar is 100 μm. (**b**) Schematic of HNO_3_ interlayer doped double layer graphene (id-DLG). (**c**) Two-photon microscopy image of pinhole defects on the graphene wires. Top and bottom wires are single- and double layer graphene, respectively. The scale bar is 10 μm. (**d**) Graphene wire resistance for single-layer graphene (SLG), double layer graphene (DLG), and interlayer doped double layer graphene (id-DLG) wires as a function of wire length. The error bars indicate the s.d. (n=4). (**e**) Optical image of high-density 256-channel graphene array. The scale bar is 1 mm.

Our id-DLG approach was effective in eliminating the defects formed in growth or fabrication. **Figure 1c** compares the graphene microwires made of single layer graphene (SLG) and id-DLG. SLG can exhibit defects formed during fabrication which may cause open circuits when patterned to form microscale wires. These defects constitute a limit in scaling graphene wire width to build high-density and large area microelectrode arrays. id-DLG overcomes this issue since these defects are randomly distributed and they do not overlap across the top and bottom layers, allowing continuous conductivity for the microscale graphene wires with high yield (**Figure 1c and Figure S2**). Furthermore, scaling down the width of long graphene wires increases the wire resistance. HNO_3_ is well known as a p-type dopant for graphene, inducing Fermi level shift in graphene due to the surface charge transfer between HNO_3_ and carbon, which increases the conductivity [65]. id-DLG formed with HNO_3_ doping effectively reduces the graphene sheet resistance from 1908 Ω/sq (SLG) to 276 Ω/sq (id-DLG) (**Figure 1d**), which enabled us to shrink the graphene wires without further attenuation in recorded signals. It is important to note that double layer graphene without interlayer dopants only reduces the sheet resistance to 606 Ω/sq (DLG).

By addressing the defect and sheet resistance issues, our id-DLG approach allowed us to fabricate transparent arrays consisting of 64 and 256 electrodes with 20 μm openings, 350 μm center-to-center pitch, and total areas of 3.1 × 2.8 mm^2^ and 6.4 × 6.1 mm^2^, respectively (**Figure 1e and Figure S3a-b**). Moreover, we designed and fabricated different configurations of transparent arrays including various opening sizes, and a dense array with 50 μm center-to-center pitch size to suit the specific needs of different in-vivo experiments (**Figure S3c-e**).

### Overcoming quantum capacitance to build ultra-small electrodes

Scaling down the electrode dimensions is important for recording high-frequency activity and building high-density arrays [61, 63]. However, ultra-small graphene electrodes exhibit large impedance due to quantum capacitance of graphene, which is a result of low density of states near the Dirac point [56]. As shown in **Figure 2a**, employing multilayer graphene and introducing dopants increase the quantum capacitance [27, 56, 65], however overall capacitance is still dominated by the quantum capacitance since it is larger than electrical double layer capacitance (see **Methods**). To reduce the impedance, we electrochemically deposited platinum nanoparticles (PtNP) which has been suggested to overcome the quantum capacitance effect by creating a low impedance parallel conductance path [56]. PtNP modifies the electrode/electrolyte interface by increasing the effective surface and enabling faradaic reactions. **Figure 2b** shows the microscope and scanning electron microscopy (SEM) images of id-DLG electrode before and after platinum nanoparticles (PtNP) deposition. The impedance distribution of an array with 64 channels before and after PtNP deposition is shown in **Figure 2c**.

**Figure 2.**
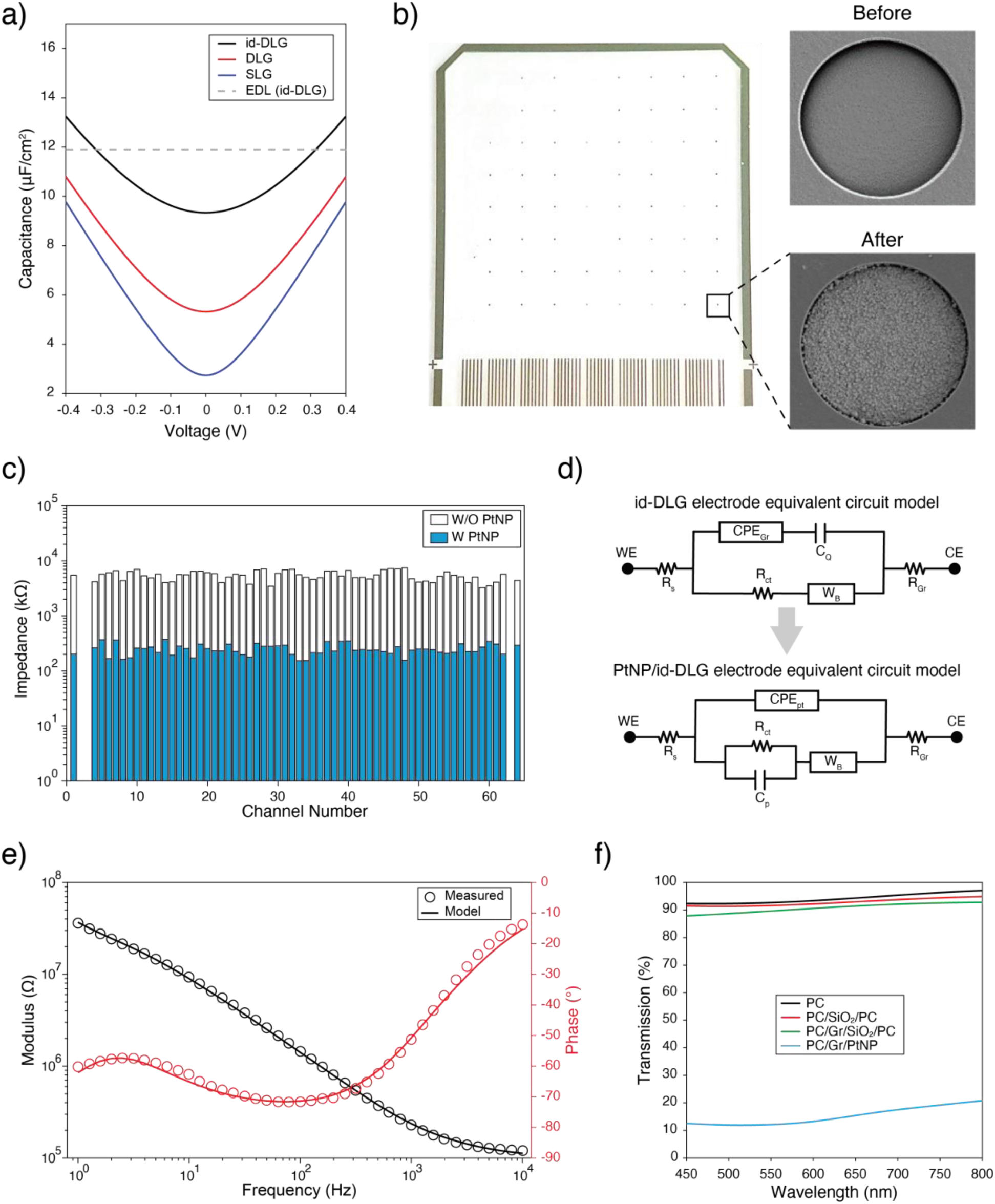
Overcoming quantum capacitance and reducing the impedance with PtNP deposition. (**a**) The quantum capacitance for single-layer graphene (SLG), double layer graphene (DLG), interlayer doped double layer graphene (id-DLG), and the Helmholtz electrical double layer (EDL) capacitance with respect to voltage are plotted. The quantum capacitance is dominant in the open-circuit potential range of graphene (-100 to 100 mV). (**b**) Optical image of the 64-channel array (left) and example SEM images of the electrode area before and after PtNP deposition (right). The scale bar is 5μm. (**c**) The impedance distribution of 64 channels at 1 kHz measured before and after PtNP deposition. The average impedances of electrodes are 5.4±1.1MΩ and 250±56kΩ (mean±s.d.), before and after PtNP deposition, respectively. (**d**) The equivalent circuit model for the id-DLG electrode with and without PtNPs. R_s_ is the solution resistance, R_Gr_ is the graphene wire resistance, C_Q_ is the quantum capacitance, CPE_Gr_ and CPE_Pt_ are the constant phase elements representing EDL of id-DLG and PtNP/id-DLG electrodes, respectively. W_B_ is the bounded Warburg element explaining diffusion process, and R_ct_ is the charge-transfer resistance that simulates Faradaic reactions. WE and CE stand for working electrode and counter electrode, respectively. (**e**) Measured electrochemical impedance spectroscopy (EIS) of the PtNP/id-DLG electrode and the fitted values using the equivalent circuit model. (**f**) Transmittance of different stacks that constitute the array. PC and Gr stand for Parylene-C and graphene, respectively.

To quantitatively analyze the electrochemical impedance of electrodes, we constructed an equivalent circuit model for id-DLG with and without PtNPs (**Figure 2d**). We modified the conventional Randles model to capture the quantum capacitance effect, resistance of graphene wires, and pseudo-capacitance of PtNP. Unlike previously reported circuit model for PtNP/SLG electrode [56], we do not have a parallel branch to explain the electrochemical reaction at the electrolyte/electrode interface as the graphene channels are completely covered by PtNPs and the interface is converted from electrolyte/id-DLG to electrolyte/PtNP. Therefore, we removed the quantum capacitance component and added C_p_ that simulates the pseudo-capacitance of PtNP. Measured electrochemical impedance spectroscopy (EIS) and the fitted equivalent circuit model results are shown in **Figure 2e**, and the extracted parameters are listed in **Table S1** for PtNP/id-DLG and id-DLG models (**Figure S4a** and see **Methods**). We observed that the impedance of channels decreased with increased deposition time (**Figure S4b)**. Since the impedance of channels saturated after 150s of PtNP deposition with a value around 200 kΩ, we decided to set the deposition time to 150s. In addition, we found that the PtNP coverage and particle grain size increased with the deposition time (**Figure S4c**). Cyclic voltammetry (CV) results before and after 150s PtNP deposition are shown in **Figure S4d**. The absence of redox peaks in the CV curve of id-DLG indicates that the electrolyte/id-DLG interface is fully capacitive. On the other hand, PtNP deposited id-DLG shows surface oxide reduction peaks around -300 mV and hydrogen absorption peaks around -500 mV to -800 mV, showing that PtNPs are contributing to the charge transfer process the electrode/electrolyte interface [56, 66]. Finally, deposition of PtNPs significantly increased the charge storage capacity (CSC) 7.5 times from 4.08 mC/cm^2^ (id-DLG) to 30.72 mC/cm^2^ (PtNP/id-DLG) due to the large pseudo-capacitance of the PtNP interface and increased surface roughness (larger effective surface area). Although the transparency of electrodes covered with PtNPs are reduced, they only cover 0.23% of the total area of the array, therefore the PtNP/id-DLG arrays maintain high transparency (**Figure 2f**). With the combination of id-DLG and PtNP we successfully achieved high-yield completely transparent arrays with ultra-small electrodes and low impedance.

### In-vivo multimodal experiments with transgenic mice

We performed multimodal experiments with transparent PtNP/id-DLG arrays to record electrophysiological signals from the cortical surface while conducting calcium imaging with two-photon microscopy from the ipsilateral visual cortex of transgenic mice expressing GCaMP6s in most cortical excitatory neurons (CaMK2-tTA::tetO-GCaMP6s; see **Methods**) [67, 68]. Drifting gratings were used as visual stimulation (**Figure 3a** and see **Methods**). Two-photon imaging was performed at two different depths, 50 μm and 225 μm, corresponding to layer 1 and layer 2/3, respectively. The field of view was 960 μm × 960 μm that spans over nine channels (**Figure 3b**). While the imaging was performed in layer 1 or layer 2/3, we simultaneously recorded the neural activities using the 64 channels of the array that spans an area of 2.5 mm × 2.5 mm. **Figure 3c** shows representative cortical potentials recorded by 64 channels during one trial of visual stimulus.

**Figure 3.**
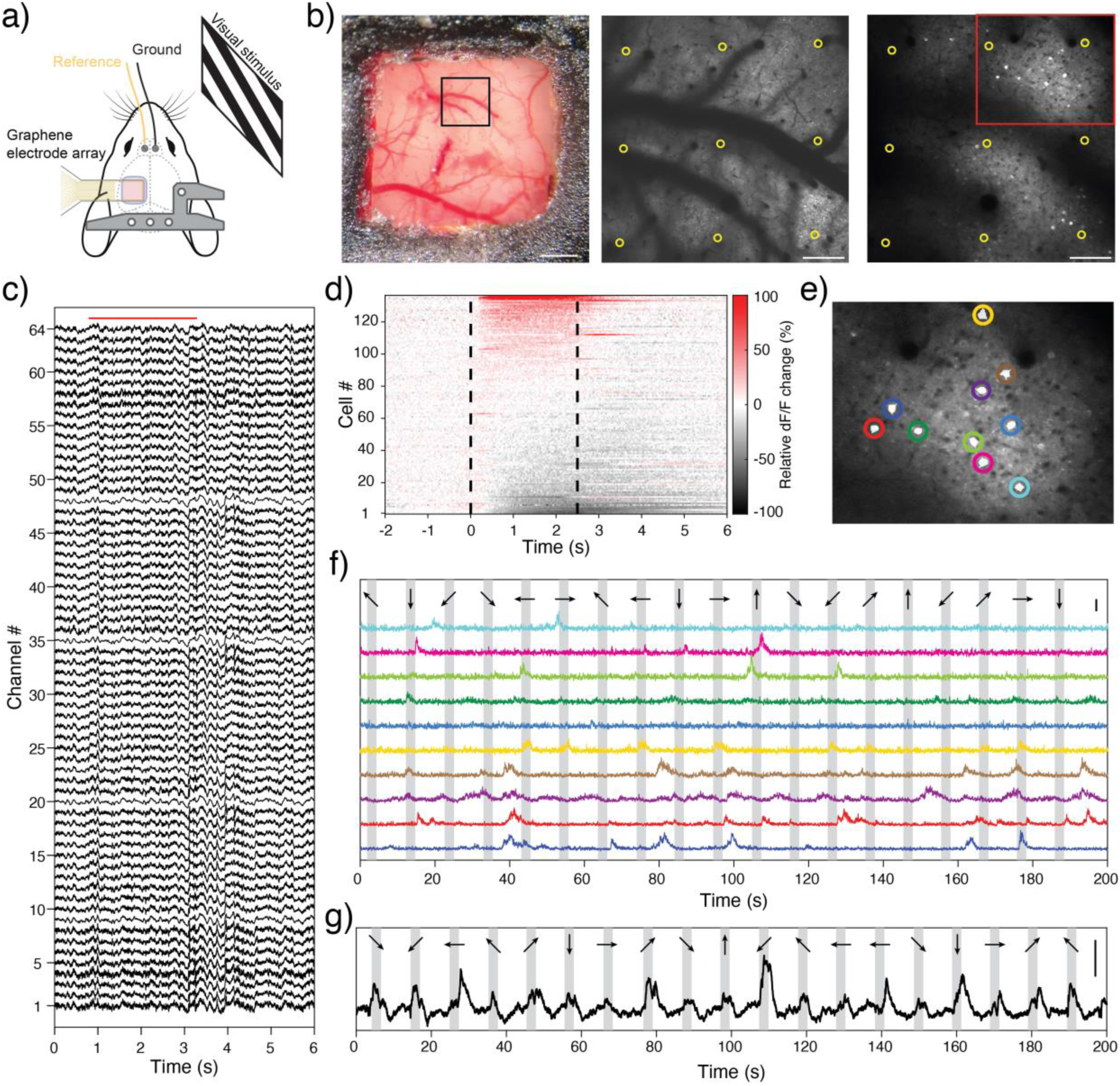
Multimodal experiments combining recordings of cortical potentials from surface and two-photon imaging at two different depths. (**a**) Schematic of the multimodal experimental setup. (**b**) Exposed cortex area covered by the array with the FoV depicted by the black square (left) and time-averaged two-photon images of layer 1 (middle) and layer 2/3 (right). PtNP/id-DLG electrodes are shown by yellow circles. The scale bars are 700 μm for the left panel and 150 μm for the middle and right panels. (**c**) Representative surface potentials recorded from the 64 channels of the array. The red line shows the duration of the visual stimulus. (**d**) Trial averaged population activity (relative to the 2-second baseline before stimulus onset) of neurons detected in layer 2/3. Black dashed lines show the onset and offset of the visual stimulus. (**e**) Ten neurons highlighted from the red box in panel b and (**f**) their normalized ΔF/F signals. (**g**) Pixel-level average ΔF/F signal of layer 1. The scale bars in panels f and g are 5 z-score. The black arrows and gray bars if panels f and g show the direction and duration of drifting gratings, respectively.

The high optical transparency of the implanted array allowed for easy detection of excitatory neurons and their compartments and recording calcium signals with single-cell resolution. The imaging quality was not compromised by the transparent graphene array and the ultra-small PtNP electrodes did not obstruct the FoV (**Figure S5**). Following motion correction and detection of neural regions of interest (ROIs), we extracted fluorescence signals and calculated ΔF/F using the Suite2p software (see **Methods**). Representative fluorescence activities of ten neurons (highlighted in **Figure 3e**) are plotted in **Figure 3f**. The trial averaged population activity shows that the imaged cells could be categorized into three groups based on their specific responses to the stimulus; activated, suppressed, and non-modulated (**Figure 3d** and see **Methods**). Activated cells exhibit an increase in their activity while suppressed cells show decreased activity during stimulus presentation. Non-modulated cells do not show significant changes in their activity during visual stimulation.

Unlike layer 2/3, layer 1 is occupied mainly by intermingled neuropils, including dendrites and axons extended from deeper layers. Therefore, the observed fluorescence represents dendritic and axonal activity. As there are almost no detectable cell bodies at this depth, the activity of layer 1 is defined as the average (pixel-level) fluorescence changes in the FoV (excluding the blood vessels, see **Methods**). **Figure 3g** shows a representative average calcium activity of layer 1 in response to drifting gratings presented in eight different orientations.

Our flexible array enabled us to record the surface potentials from 64 channels that spanned over a large area (2.5 X 2.5 mm) of the cortex including regions such as primary visual cortex (V1), primary somatosensory cortex (S1), posterior parietal cortex (PPC), and retrosplenial cortex (RSC) (**Figure 4a**). With such broad spatial coverage, we were able to examine the propagation of visual stimulation responses. The responses were initiated from the top-right parts of the array that are located over the visual cortex and spread to the other channels while the peak amplitudes are detected over V1 (**Figure 4b**). The top three rows of the array, which were placed over V1 and RSC, appeared to have the strongest biphasic responses. We analyzed the power of visual evoked responses at different frequency bands and found that the high-frequency bands (γ, and MUA) were more localized compared to low-frequency bands (δ, θ), which propagated to RSC, PPC, and even S1 (**Figure 4c** and see **Methods**). This result is consistent with previous works that showed the spatial reach of signal is limited at higher frequencies [62, 69–71]. The small electrode size (20 μm) with low impedance allowed us to record multi-unit activity (MUA) from the cortical surface with high fidelity. Representative MUA traces detected on different channels are shown in **Figure 4d**. These short-duration spikes recorded from the surface were classified as MUA since their auto correlograms do not show any refractory period (**Figure S6a**). To investigate the origins of MUA spikes detected from the surface, we examined the correlation between the cellular signals from calcium imaging and the MUA power for each channel. To calculate the MUA power, we band-pass filtered the signal between 500 Hz to 4 kHz and smoothed the squared values with a Gaussian kernel (see **Methods**). First, we extracted the peaks of the cell-averaged calcium signal and then took the time-average of MUA power around those peaks’ onsets for all 64 channels (see **Methods**). The high correlation between the cellular calcium peaks and the MUA for the channels within the FoV (**Figure 4e**) suggests that the spiking activity of L2/3 excitatory neurons underneath these channels is an important contributor to the MUA signals detected on the surface. **Figure 4f** shows representative cell-averaged calcium signal and MUA power of the channel with maximum correlation. The correspondence between the two signals is evident from the sharp deflections in the MUA power followed by peaks in the calcium signal. **Figure 4g** shows the correlation between calcium peaks and MUA power extracted from the whole recording of the same channel. We found similar correlation values between MUA power and calcium signal in other experiments with 16ch PtNP/id-DLG arrays (**Figure S6b-e**).

**Figure 4.**
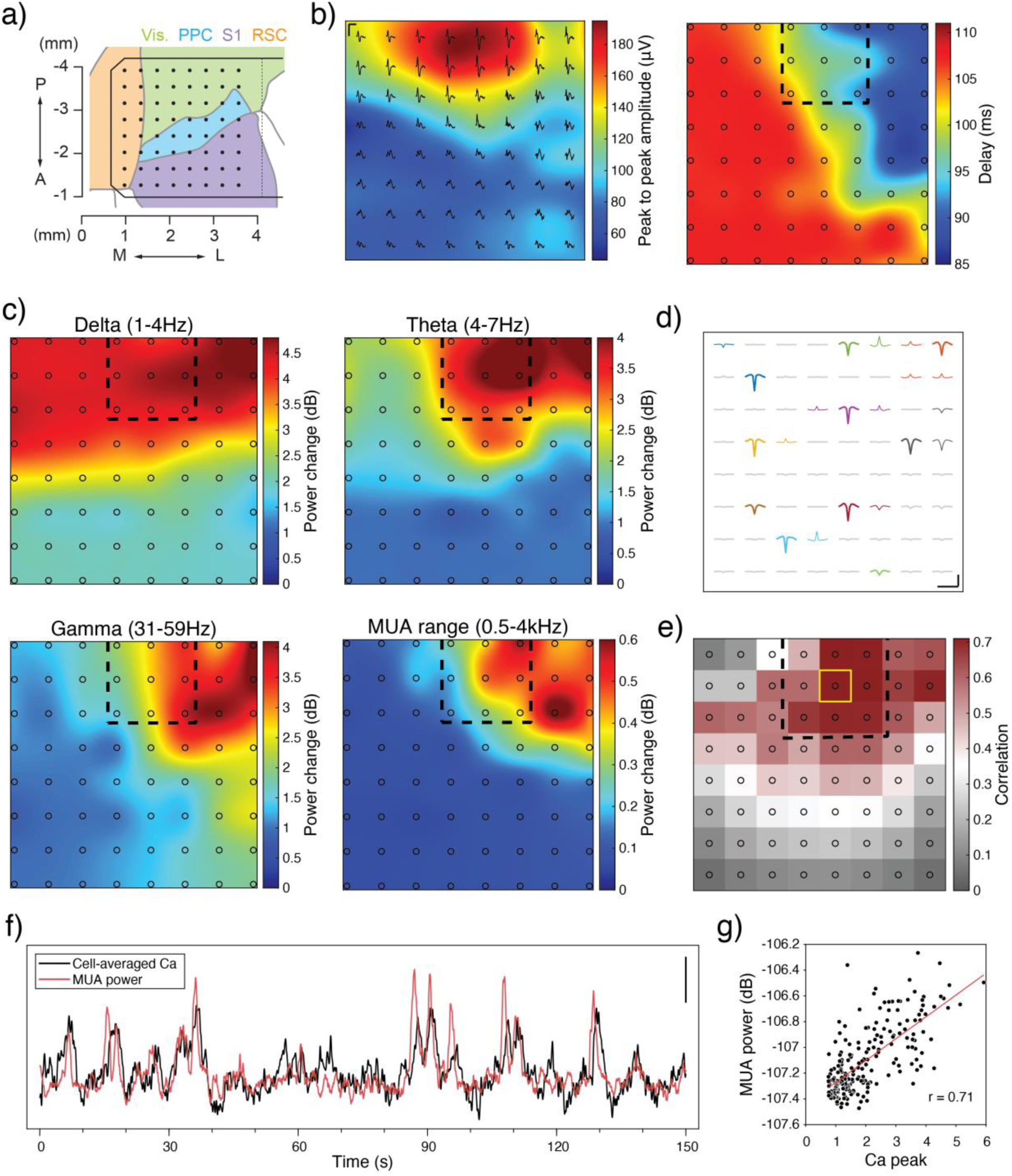
Stimulus-evoked local field potentials and high-frequency activities detected using electrodes on the cortical surface. (**a**) Cortical regions covered by the 64 channels of the array. The total area covered is 2.45 X 2.45 mm. (**b**) Peak-to-peak amplitude (left) and the delay map (right) of the visual evoked responses. The horizontal and vertical scale bars are 250 ms and 100 μV, respectively. (**c**) Spatial maps of the evoked powers (relative to the baseline) at different frequency bands across the array. High-frequency activities are spatially localized while low-frequency bands have broad propagation ranges. (**d**) Representative event-triggered MUA waveforms on different channels. The horizontal and vertical scale bars are 2 ms and 20 μV, respectively. (**e**) Correlation between the cell-averaged calcium peaks and MUA power around the peak onset for all 64 channels. The channels in the FoV show the highest correlation values. Yellow box shows the channel with maximum correlation (r=0.71). Black dashed boxes and black circles in the colormaps in panels b, c, and e indicate the FoV and the electrodes’ locations, respectively. (**f**) Representative cell-averaged ΔF/F and MUA power of the channel with maximum correlation (yellow box in panel e). The scale bar is 2 z-score for calcium and 0.5 dB for MUA power. (**g**) Scatter plot of cell-averaged calcium peaks and corresponding MUA powers for the channel with maximum correlation (yellow box in panel e).

### Predicting Neural Activity in Layer 1 and Layer 2-3 from Surface Recordings

Given the correlation between the MUA power recorded from the surface and the cellular calcium signals imaged at 225 μm depth, we asked whether it is possible to predict the brain activity at deeper layers by only harnessing high-resolution electrical recordings from the cortical surface. To that end, we implemented a simple neural network model that consists of a linear hidden layer, a single-layer LSTM network, and a linear readout layer (**Figure 5a**) [22, 72]. The neural networks were trained to learn the nonlinear relationships between cellular calcium activities and surface potentials. It is important to emphasize that simultaneous recordings enabled by the high transparency of our graphene microelectrode arrays are critical to acquiring the multimodal datasets needed for training the neural networks. The power of signals at different frequency bands (δ: 1–4 Hz, θ: 4–7 Hz, α: 8–15 Hz, β: 15–30 Hz, γ: 31–59 Hz, H-γ: 61–200 Hz, MUA: 0.5–4 kHz) were fed as inputs to the network to predict the pixel-level averaged calcium fluorescence change of L1 and L2/3 and the cell-averaged activity of L2/3. Five-fold cross-validation was performed by splitting the 40-minute recording sessions into eight-minute-long segments. Representative examples of decoded and ground truth activities for L1 and L2/3 are shown in **Figure 5b**. Calcium activity predicted from the surface potentials shows good agreement with the ground truth calcium fluorescence change imaged using two-photon microscopy for both depths. To evaluate the contributions spatially provided by different channels, we performed the decoding using subsets of channels starting from those closest to the FoV (**Figure S7a**). The decoding performance increased with the inclusion of more channels (**Figure 5c**), which indicates that different channels provide complementary information. However, the decoding performance was saturated when ∼20 channels were used, suggesting that additional channels provide redundant information beyond this point.

**Figure 5.**
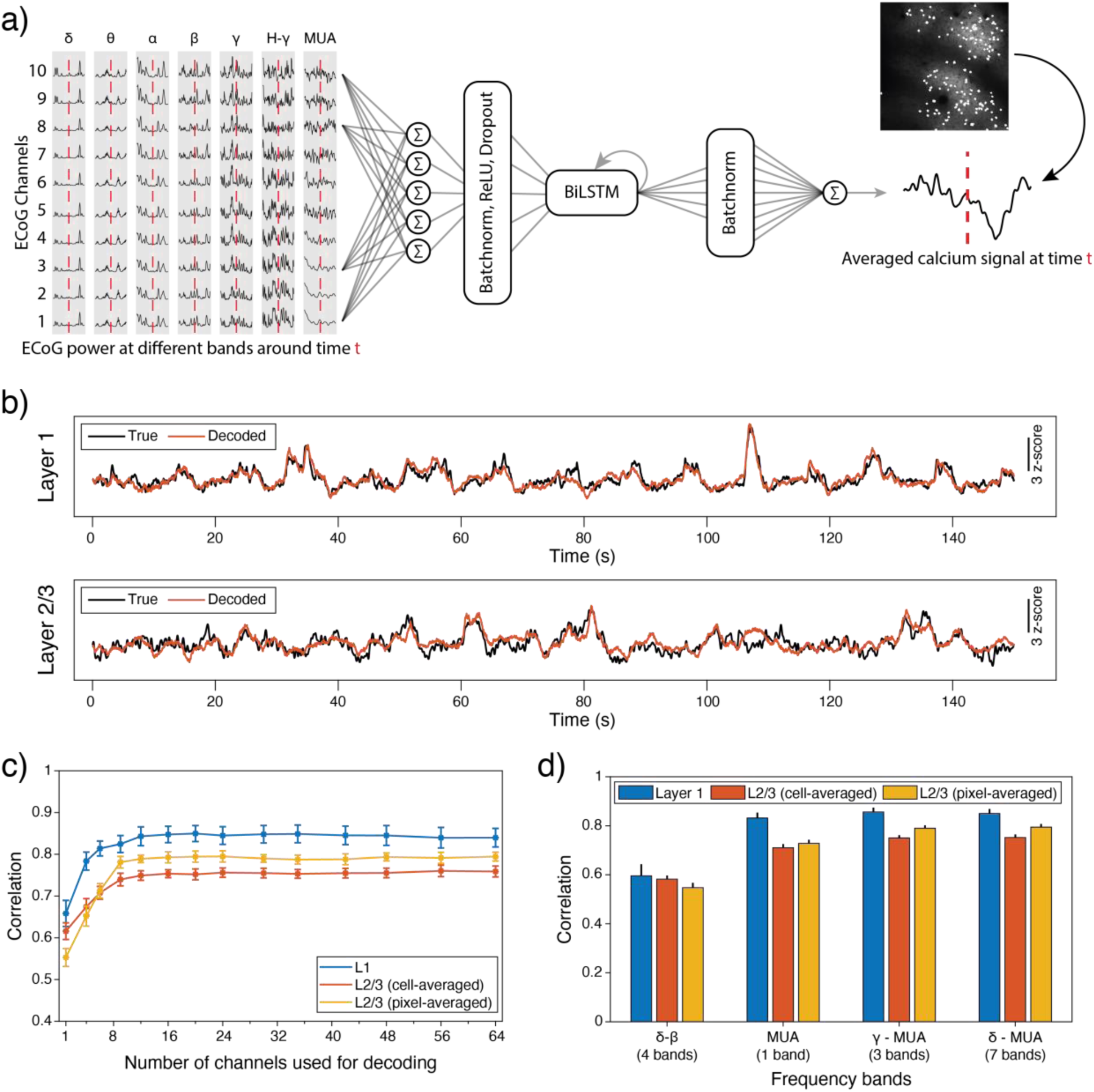
Decoding the average calcium activity from recorded surface potentials. (**a**) Schematic of the decoding model. Signal powers at different frequency bands (ten channels are shown as example) around time t are used as inputs to the model to decode the calcium activity at time t. The model consists of a linear hidden layer, a single-layer BiLSTM network, and a linear readout layer. (**b**) Decoded (orange) vs ground truth (black) ΔF/F of layer 1 (pixel-averaged) and layer 2/3 (cell-averaged). (**c**) Decoding performance for layer 1 and layer 2/3 (cell and pixel-averaged) using all seven frequency bands but different numbers of channels. The error bars indicate the s.e.m. (**d**) Decoding performance for layer 1 and layer 2/3 (cell and pixel-averaged) using different frequency bands of the 20 channels closest to the FoV. Bars and black lines indicate the mean and s.e.m., respectively.

We next investigated the contribution of different frequency bands to the decoding performance by carrying out decoding using low (δ, θ, α, and β) and high (γ, H-γ, MUA) frequency components from 20 channels closest to the FoV (9 channels in the FoV and 11 channels around it, see **Figure S7a**). The results show that the highest correlation is achieved when MUA and γ, H-γ were included, suggesting that high-frequency components carry a vast amount of information on the neural activity including the cellular spiking in the FoV (**Figure 5d**). As demonstrated in **Figure 4c**, the low-frequency bands were also modulated by the visual stimulation, so the model could still use these bands as informative features to decode the calcium activity. However, the correlation between the peaks of average cellular activity and power at different frequency bands for those channels over and around the FoV (**Figure S7b** and see **Methods**) is significantly larger for high frequency bands (H-γ and MUA). Therefore, excluding the low-frequency components does not have a substantial effect on the decoding performance of cell averaged calcium activity. This indicates that low-frequency components do not provide additional information when combined with high-frequency bands for decoding cellular spiking at deeper layers.

### Predicting Single-cell Activity from Surface Recordings

We showed that recordings from cortical surfaces could be used to train networks and predict the average calcium fluorescence change of neurons in L2/3. However, the average calcium signal mostly represents the dominant and synchronous dynamics in the neural network. A more interesting question is whether predicting calcium fluorescence of single cells from deeper layers is possible by only using high-resolution recordings of cortical potentials. Developing a network similar to **Figure 5a** to predict the activity of all 136 neurons would require increasing the complexity of the network which is not efficient due to the covariances in the neural activity. Previous studies have shown that the neural activity of neurons could be defined by low-dimensional manifolds that capture most of the variance [73, 74]. Therefore, a better approach would be predicting the low-dimensional neural manifolds and projecting them back to the single-cell space. Gaussian Process Factor Analysis (GPFA) is a generative model that unifies dimensionality reduction and smoothing in one framework to extract latent representations that describe the shared variability of high-dimensional data [75].

To investigate the feasibility of predicting the single-cell activities of L2/3 neurons from the surface potentials, we first used GPFA to find a low-dimensional latent space that is very representative of the high-dimensional calcium fluorescence signal. We identified eight distinct latent variables that explain most of the variance of the high-dimensional data (**Figure S8a**). Next, we trained eight networks (same architecture used for the average calcium fluorescence decoding) to predict each of these latent variables separately using the surface recordings of 20 channels closest to the FoV (**Figure S8b**). To reconstruct the single-cell calcium fluorescence, we projected the decoded latent variables to high dimensional space using the GPFA parameters. The schematic in **Figure 6a** shows the three main steps in single-cell decoding which are dimensionality reduction using GPFA, prediction using RNNs, and projection to high dimensional space (See **Methods** and **Figure S8c**). Representative examples of decoded and ground truth single-cell activities are shown in **Figure 6b**, which demonstrates that our model can infer the activity of several neurons at depth using electrical recordings from the cortical surface. **Figure 6c** shows the decoding performance of all 136 cells with their location in the FoV. It is noteworthy to mention that maximum correlation is partially limited by the amount of information extracted using the GPFA model, as seen in the reconstructed calcium signals using true latent variables (**Figure S9a-b**). Prediction error can be further reduced, and correlation values can be further increased by optimizing the dimensionality reduction methods. We also compared the decoding performance for modulated (suppressed or activated) and non-modulated cells and found that the decoding performance is significantly better for cells that are responsive to the visual stimulus (**Figure S9c,** see **Methods**). These results indicate that the surface potentials recorded by transparent PtNP/id-DLG arrays carry information about the neural activities in both superficial and deep layers of the brain and could be used to infer neural population dynamics not only at the average but even at single-cell level by projecting higher dimensional neural activity to a latent space.

**Figure 6.**
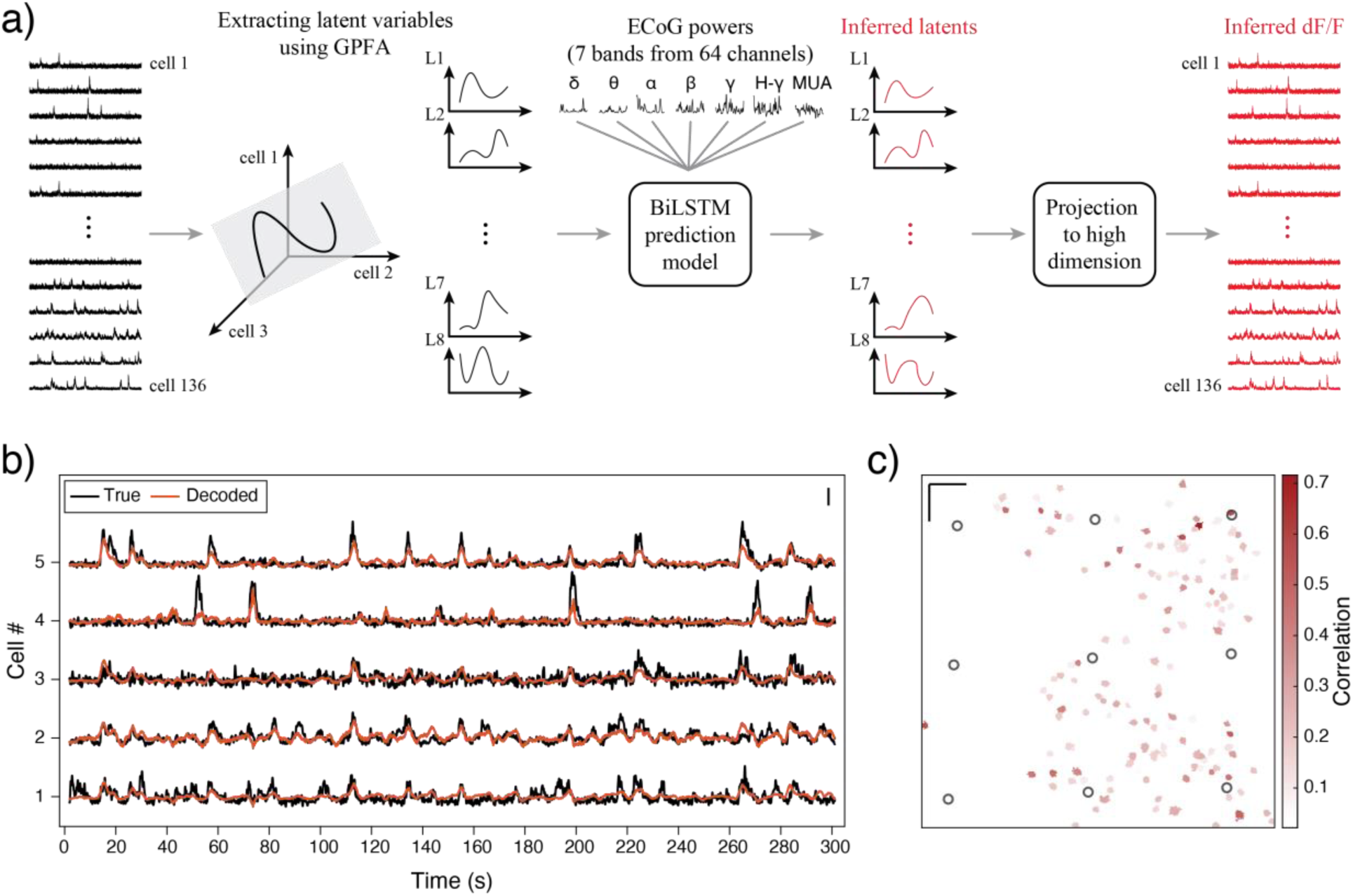
Decoding single-cell calcium activity from surface potentials using latent variables. (**a**) Schematic of single-cell decoding model. Eight latent variables (L1 to L8) extracted using GPFA are used to train BiLSTM models (similar to Fig. 5a). Inferred latent variables are projected to high-dimensional space to achieve single-cell ΔF/F signals. (**b**) Representative examples for decoded (orange) vs ground truth (black) ΔF/F of five best-decoded cells. The scale bar is 3 z-score. (**c**) Decoding performance for all 136 cells with their locations outlined in the FoV. Black circles are the 9 channels inside the FoV. The scale bars are 100µm.

## Discussion

In this work, we developed a transparent, high-density graphene array with ultra-small electrodes and demonstrated its application in multimodal experiments to study the neural dynamics at different cortical layers with complementary spatiotemporal resolution provided by optical imaging and electrophysiological recording. Complete transparency of the graphene arrays enabled us to perform multimodal experiments combining electrical recordings from surface with two-photon imaging from depth and investigate the neural dynamics in the visual cortex of awake mice presented with drifting gratings as visual stimuli. By using double layer graphene, interlayer doping, and PtNP deposition we achieved high-aspect-ratio graphene wires and ultra-small electrodes with low impedances and drastically reduced the artifacts induced by two-photon imaging.

We explored the multimodal datasets and found that the trial averaged signal powers at different frequency bands (δ, θ, γ, and MUA) demonstrated different spatial propagation patterns. Consistent with previous studies on the propagation of signals, we found that the responses are more localized at higher frequency bands. We also realized that the peaks of cell-averaged calcium activity of L2/3 neurons are highly correlated with the increases in the MUA power of those channels in and around the FoV. Such correspondence indicates that the synchronous spiking activity of pyramidal cells propagates to the cortical surfaces and could be detected using Electrocorticography (ECoG) electrodes. The means of propagation could be the axons projected to the superficial layers, volume conduction, and other connections in the neural network. These results suggest that surface potentials convey information about the neuronal activity at depth and proper features extracted from these signals could be used to predict the cellular activities at depth.

To that end, we first focused on the average calcium signals in L1 and L2/3. Due to the absence of neuronal bodies in L1, we used pixel-averaged ΔF/F as representative output signal and for L2/3 we used both the cell- and pixel-averaged ΔF/F signals. We developed simple RNNs and fed the powers of 7 different frequency bands (δ, θ, α, β, γ, Hγ, and MUA) as predictive features to decode the average calcium activities in L1 and L2/3. The inferred calcium signals resemble the true signals with minimal error which demonstrates the performance of the decoding model. To investigate the spatial and frequency contribution of these features we repeated the decoding with different combinations of channels and frequency bands. We showed that the inclusion of more channels improves the decoding performance due to the non-redundant information provided by each channel. However, adding the channels that are farther away from the FoV does not improve the decoding performance which suggests that these channels do not provide additional information about the activity of imaged neurons. We also showed that excluding the low frequency (δ, θ, α, β) signals does not deteriorate the decoding performance which suggests that these bands do not provide additional information when combined with high frequency bands (γ, Hγ, and MUA).

To further improve the spatial resolution of the decoding network, we used GPFA to extract a low-dimensional neural manifold and trained RNNs to separately decode the extracted latent variables. By projecting the decoded latent variables, we reconstructed single-cell calcium activities of all 136 neurons in L2/3. We predicted the activity of several neurons with high correlation (r>0.5) which indicates that surface recordings convey information on the spiking activity of neurons at depth.

These results demonstrate that our transparent graphene arrays could be potentially integrated with other techniques to facilitate multimodal experiments with unprecedented spatiotemporal resolutions. For instance, optical techniques could be utilized to manipulate/monitor the neural circuits and uncover the complex dynamics of surface potentials by realizing cross modality inference. Ultimately this may lead to localizing the potential sources of distinct features that are detected in surface recordings. The results of such experiments could be the applied to BCI technologies to improve and expand current systems to new realms of complex motor and behavioral tasks. Moreover, such multimodal experiments could be used to study the generation and propagation of neural oscillations that are imperative to various cognition mechanisms. Recordings of neural activity at depth without implanting invasive neural probes could extend the lifetime of neural implants and improve the longevity of BCI technologies and pave the way for their medical translation. It can also open up new avenues for minimally invasive neural prosthetics or targeted treatments for various neurological disorders.

## Methods

### Fabrication process of id-DLG arrays

To form the transparent and flexible substrate, we deposited a 14 µm-thick layer of Parylene-C on a 4-inch silicon wafer coated with 100 nm PMGI SF3 as sacrificial layer. Next, we DC sputtered 5 nm Chromium and 100 nm gold on the parylene-C substrate and patterned it with photolithography and wet etching to form metal wires and contact pad. The First graphene layer was transferred using electrochemical delamination process previously developed [26, 76]. To decrease the wire resistance, the first graphene layer was immersed into 50% nitric acid (HNO_3_) solution for 10 minutes. After cleaning the HNO_3_-doped graphene with acetone and IPA, the second graphene layer was transferred using the same process as the first layer. To pattern double layer graphene, we used bilayer photoresist (PMGI/AZ1512) and etched the graphene with oxygen plasma, followed by acetone/IPA cleaning. To protect the double layer graphene during the next steps, we sputtered a 25 nm silicon dioxide (SiO_2_) etch-stop layer on the patterned graphene. Then we deposited a 2 µm-thick Parylene-C as encapsulation layer and patterned it with oxygen plasma to define electrode openings. To remove the protective SiO_2_ layer and get access to the double layer graphene we used 6:1 buffered oxide etchant. Finally, we detached the arrays from the wafer by immersing it in acetone and applying slight physical force to the edges of the wafer.

### Electrical double layer capacitance and quantum capacitance calculation

To extract the capacitances, we first obtained the values of CPE_Gr_ and C_Q_ by fitting the measurement data to the circuit model of id-DLG electrode. Then we extracted the CPE parameters (capacitance parameter, Y, and phase change element exponent, α) and used equation (1) to calculate the C_dl_ [77]. R_s_ is the solution resistance.

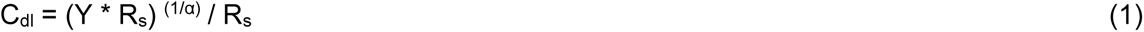

We used equation (2) and the measured open circuit voltage to calculate the impurity concentration of SLG, DLG, and id-DLG [64, 78].

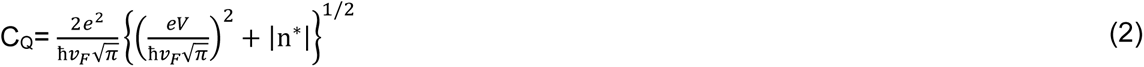

Here, *v*_F_ is Fermi velocity, ħ is plank constant, and V is open circuit voltage. To plot the capacitances in **Figure 2a**, we used equation (2) and swept the open circuit voltage from -0.4 to 0.4 V (Cdl is not a function of V, so its value is constant).

### Electrode characterization and platinum nanoparticle deposition

All the electrochemical characterizations were conducted with Gamry 600 plus with immersing in 1x phosphate buffered saline. Both electrochemical impedance spectroscopy and cyclic voltammetry were measured under three-electrode configuration using Ag/AgCl as reference electrode and Pt as counter electrode. To avoid electromagnetic noise, all the measurements were conducted inside of Faraday cage. Platinum nanoparticles (PtNPs) deposition was conducted with two-electrode configuration (Gamry 600 plus). The id-DLG electrode was connected to the working electrode while Pt wire was connected to the counter or auxiliary electrode. While both electrodes were immersed in the H_2_PtCl_6_ (0.05 M) and K_2_HPO_4_ (0.01 M) solution, the current of 50 nA was flown from the id-DLG electrode to counter electrode for selected time periods under ambient condition.

### Animal Procedures

All procedures were performed in accordance with protocols approved by the UCSD Institutional Animal Care and Use Committee and guidelines of the National Institute of Health. Adult mice (cross between CaMKIIa-tTA (JAX 003010) [68]) and tetO-GCaMP6s (JAX 024742) [67], 2 months old) were anesthetized with isoflurane (3% for induction and 1% for maintenance). Both eyes were protected by Vaseline (Vaseline) and a circular piece of scalp was removed. After cleaning the underlying bone using a razor blade, a custom-built head plate was implanted to the exposed skull (≈1 mm posterior to lambda) with cyanoacrylate glue and cemented with dental acrylic (Lang Dental). Two stainless steel screw (F000CE156, J.I. Morris) were implanted over olfactory bulb as reference and ground. A square craniotomy was made over the left hemisphere (∼3.5 × 4 mm, centered ∼1.75 mm lateral and 2 mm posterior to bregma), and dura of the craniotomized area was carefully removed with a hooked needle. The transparent PtNPs/id-DLG electrode array was first attached to a glass window with UV glue and connected to the amplifier board. Then the assembled interface was gently placed onto the exposed cortex with the electrode array facing to the cortical surface. The glass window was gradually press down through a micromanipulator (Sutter Instrument) until the whole electrode array was tightly attached to the cortical surface. Cortical areas covered by the electrode array included primary somatosensory cortex (S1), posterior parietal cortex (PPC), primary visual cortex (V1), and retrosplenial cortex (RSC). Vetbond (3M) was applied to fill the gap between the skull and the glass window, and the glass window was further secured with cyanoacrylate glue and dental acrylic. A cocktail of dexamethasone (2 mg/kg body weight), buprenorphine (0.1 mg/kg body weight), and baytril (10 mg/kg body weight) was given at the end of surgery. The animal was fully recovered from anesthesia before recording.

### Visual Stimulation

Square-wave drifting grating stimuli (100% contrast, 0.04 cycles/degree, 3 cycles/sec, covering entire contralateral receptive field) were presented on an LCD monitor (30 × 38 cm) positioned 15 cm away from the right eye using Psychtoolbox (http://psychtoolbox.org/). One of 8 orientations (45° apart) was presented for 2 or 2.5 sec on each trial in pseudorandom order. Inter-stimulus-interval was 8 seconds. We presented each orientation at least 30 times in a session.

### Two Photon Imaging and analysis of imaging data

Two-photon imaging was conducted for a head-fixed awake mouse through a 16 × 0.8 NA objective (Nikon) mounted on a commercial two-photon microscope (B-scope, Thorlabs) and using a 925 nm laser (Ti:sapphire laser, Newport). Images were acquired at ∼29 Hz and a resolution of 512 × 512 pixels, covering 960 × 960 µm (**Figures 2b-C**). The laser power was ∼15mW for imaging layer 1 (∼50 μm deep) and ∼40mW for imaging layer 2/3 (∼225 μm deep). Acquired images were motion corrected offline [79]. For quantification of calcium signals from layer 1, pixels in blood vessels and 10 pixels close to frame edges were excluded. The fluorescence time course (F) was calculated as the ground average of remaining pixels in each frame. At each time point, the baseline (F_0_) was estimated by the 10^th^ percentile of the fluorescence distribution. For quantification of calcium signals from layer 2/3 cell bodies, ROIs were first identified by Suite2P package [80] and then visually inspected to remove non-somatic ones. Next, fluorescence time course of each cellular ROI and its surrounding neuropil ROI was extracted using Suite2P package. Then fluorescence signal of a cell body was estimated with F_cellbody_ = F_cellROI_ − 0.7*F_neuropilROI_. ΔF/F_0_ was computed as (F_cellbody_ − F_0_)/F_0_, where F_0_ is the 8^th^ percentile of the intensity distribution during the recording session.

To analyze the stimulus response of imaged cells, we subtracted the baseline activity (2 seconds before the stimulus onset) from the trial averaged fluorescence signal for each cell body and normalized with the baseline activity. To categorize the cells, we sorted (descending) them based on their average of normalized stimulus response (between 0.3 to 3 s after the stimulus onset). We considered the first and last 20 cells as being responsive to the visual stimulation.

### Electrophysiology Data Recording

Electrophysiological recording was conducted with the RHD2000 amplifier board and RHD2000 evaluation system (Intan Technologies). The sampling rate was set to 20 kHz and DC offset was removed with the recording system’s built-in filtering above 0.1 Hz. Intan data was imported into MATLAB (MathWorks) and analyzed using custom scripts.

### Electrophysiology Data Analysis and Statistics

Analyzing the data is done mainly in MATLAB v2019b and the decoding is done using Python. Electrodes with impedances above 10MΩ are excluded from analysis. To remove common artifacts (imaging and power line), a bank of notch filters is applied to the raw surface recordings (the filters are optimized for each channel separately). The multi-unit activity (MUA) is extracted by applying a 6^th^ order bandpass filter from 0.5 to 4 kHz followed by common average referencing. The signals are lowpass filtered below 250 Hz using a 4^th^ order Butterworth filter to achieve the local field potential (LFP). The peaks of the visually evoked LFP were extracted and the trial averaged peak-to-peak amplitude and the propagation of the stimulus responses were visualized using 2-D color maps. To further filter the signals into common low frequency bands (δ: 1–4 Hz, θ: 4–7 Hz, α: 8–15 Hz, β: 15–30 Hz, γ: 31–59 Hz, H-γ: 61–200 Hz), 6^th^ order Butterworth bandpass signals are applied with corresponding frequency ranges.

The powers at different bands (δ, θ, α, β, γ, H-γ, MUA) were calculated by taking the square of the bandpass filtered signals and applying a 100ms Gaussian filter to reduce the noise. The power changes due to the visual stimulus were calculated by trial averaging the powers at different bands and subtracting the baseline activity (2 seconds before the stimulus onset) and the results were demonstrated using 2D spatial maps to visualize the localization of different bands. The MUA events for each channel were extracted by using the threshold crossing method (-4*std).

To analyze the MUA and average cellular calcium correlation, the peaks of normalized cell-averaged ΔF/F are determined (*findpeaks*, minimum peak height is set to 0.75) and the MUA power of each channel is averaged in a 2-second window [-1.5s, 0.5s] around the calcium peak onset. The Pearson correlation values were calculated for each channel between the calcium peaks and the averaged MUA powers. The same procedure is followed to calculate the correlation of average cellular calcium activity with other frequency bands (δ, θ, α, β, γ, H-γ).

To decode the cell averaged calcium activity from surface potentials, a neural network model with a sequential stack of a linear hidden layer, one bidirectional LSTM layer and a linear readout layer was implemented. The linear hidden layer is followed by batch normalization, ReLU activation, and dropout (p=0.3). The LSTM layer is followed by batch normalization. ECoG power at different frequency bands (δ, θ, α, β, γ, H-γ, MUA) were down sampled to match the sampling rates of the calcium signal (29 Hz) and clipped with a threshold of 95 percentile to suppress the potential artifacts. These power signals are then used as inputs to the neural network model. To decode the neural activity at time step t, the power segments between [t − 1.5 s, t + 1.5 s] was used (90 time-steps in total). The 1st linear layer had 25 neurons and the bidirectional LSTM had 15 hidden neurons. The last layer outputs the predicted cell-averaged calcium signal.

Adam is used to train and optimize the parameters of the model with learning rate = 6 × 10−5, beta1 = 0.9, beta2 = 0.999, epsilon = 1 × 10−8. The batch size was set to 128 and the training converged within ∼ 20 epochs. The mean squared error (MSE) was used as the loss function. Five-fold cross-validation is performed by splitting the 40 minutes recording sessions into five segments, each lasting for 8 minutes. The Pearson correlation between the decoded and ground truth data was used to evaluate the model performance, and the correlation values are averaged over five folds to get a single correlation value.

### Low Dimensional Latent Space of Population Activity

GPFA models observations as a Gaussian model that is related to the latent variable through equation (3).

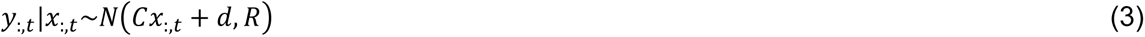

Where x_:,t_ represents the latent variable at timepoint t, d is the signal mean, C is the factor loading matrix, and R represents the covariance matrix. The ith latent variable x is modeled as a Gaussian process (GP) with a covariance matrix K that correlates latent variables across time points:

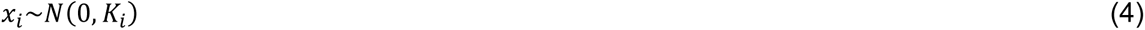

Using the training data of calcium signal Y, we train a GPFA model that learn the parameters and infer the trajectory of the latent variable x.

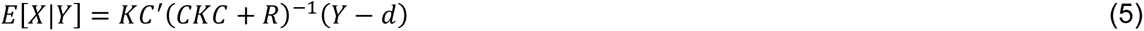

We use the latent variables *x* and the inferred latent variable 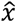 from the BiLSTM model to project into calcium signal space **(Fig S8c)**.

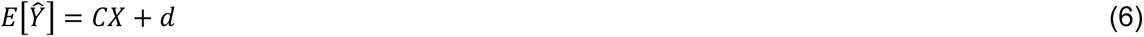

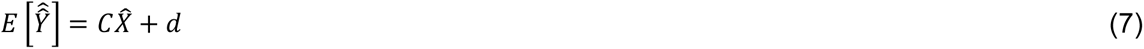

where 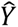 is the projected calcium signal using the originally inferred latents and 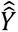 is the projected calcium signal using the predicted latents from the BiLSTM model. The projected calcium signals are then compared to the true calcium signals **(Fig S9)**.

## Acknowledgments

We thank members of the Kuzum, Komiyama, and Cubukcu labs for helpful discussions. This research was supported by grants from the ONR Young Investigator Award (N00014161253), NSF (ECCS-2024776, ECCS-1752241 and ECCS-1734940) and NIH (R21 EY029466 and R21 EB026180 and DP2 EB030992) to D.K., and grants from NIH (R01 NS091010A, R01 EY025349, R01 DC014690, R21 NS109722, and P30 EY022589), NSF (1940202), Pew Charitable Trusts, and David and Lucile Packard Foundation to T.K. Fabrication of the electrodes was performed at the San Diego Nanotechnology Infrastructure (SDNI) of UCSD, a member of the National Nanotechnology Coordinated Infrastructure, which is supported by the National Science Foundation (Grant ECCS-1542148).

## Author Contributions

This work was conceived by M.R., J.-H.K., and D.K. Microelectrode array fabrication was performed by J.-H.K and the characterization of arrays was performed by J.-H.K, C.D.-E., and M.W. All animal experiments were performed by C.R., X.L. and M.R. and analyzed by M.R. with contributions from J.-H.K., A.A., V.G., and D K. The manuscript was written by M.R., J.-H.K., and D.K. and edited by all authors.

## Data availability

The data that support the findings of this study are available from the corresponding authors upon reasonable request.

## Code availability

The code for processing the neural data and decoding is available from the corresponding authors upon request.

